# Rainforest-to-pasture conversion stimulates soil methanogenesis across the Brazilian Amazon

**DOI:** 10.1101/2020.03.08.982587

**Authors:** Marie E. Kroeger, Laura K. Meredith, Kyle M. Meyer, Kevin D. Webster, Plinio Barbosa de Camargo, Leandro Fonseca de Souza, Siu Mui Tsai, Joost van Haren, Scott Saleska, Brendan J.M. Bohannan, Jorge L.M. Rodrigues, Klaus Nüsslein

## Abstract

The Amazon rainforest is a biodiversity hotspot and large terrestrial carbon sink that is threatened by agricultural conversion. Rainforest-to-pasture conversion leads to the release of a potent greenhouse gas by converting soil from a methane sink into a source. The biotic methane cycle is driven by microorganisms; therefore, this study focused on active methane-cycling microorganisms and their functions across land-use types. We collected intact soil cores from three land use types (primary rainforest, pasture, and secondary rainforest) of two geographically distinct areas of the Brazilian Amazon (Santarém, Pará and Ariquemes, Rondônia) and performed DNA stable-isotope probing coupled with metagenomics to identify the active methanotrophs and methanogens. At both locations, we observed a significant change in the composition of the isotope-labeled methane-cycling microbial community across land use types, specifically an increase in the abundance and diversity of active methanogens in pastures. We conclude that a significant increase in the abundance and activity of methanogens in pasture soils could explain the greater methane flux. Furthermore, we found that secondary rainforests recovered as methane sinks, indicating the potential for reforestation to offset greenhouse gas emissions in the tropics. These findings are critical for informing land management practices and global tropical rainforest conservation.

## INTRODUCTION

Climate change, caused by the anthropogenic release of greenhouse gases^1^, is affecting every ecosystem on Earth. Although the majority of greenhouse gases released to the atmosphere are associated with the industrial revolution and fossil fuel combustion, land use change is a significant contributor. Specifically, tropical deforestation in the last decade has released ∼1 Pg C yr^−1^, an equivalent to 10% of anthropogenic carbon dioxide emissions^1^, and 78% of total greenhouse gas emissions in Brazil are caused by land use change^2–3^. In addition to being biodiversity hotspots of plants and animals, tropical rainforests are large terrestrial carbon sinks. However, rainforest deforestation to create cattle pastures or agricultural fields releases large amounts of stored carbon, converting former terrestrial carbon sinks into major carbon sources^3–4^. In the Amazon rainforest particularly, over 1 Mha of forest has been lost in 2017 alone^5^. The conversion of primary rainforest (i.e. mature rainforest older than 150 years) to cattle pasture is a main cause of deforestation in Brazil and not only changes plant diversity but also the microorganisms that drive soil biogeochemical cycling^6^.

The methane (CH_4_) biogeochemical cycle is of interest because of its potency as a greenhouse gas with 86-times the global warming potential of carbon dioxide over a 20-year timescale^1^. Biotic CH_4_ cycling is controlled by microorganisms, specifically methanogenic archaea that produce CH_4_, and methanotrophic bacteria that consume CH_4_^7–8^. The balance between these two functional groups determines whether the soil acts as a CH_4_ source or sink. Under anoxic conditions, soil methanogenic archaea generally metabolize fermentation products such as acetate (acetoclastic methanogenesis) or reduce carbon dioxide with hydrogen (hydrogenotrophic methanogenesis) to produce CH_4_^9–11^. Methanotrophs are commonly aerobic bacteria located at anoxic/oxic boundaries from either *Gammaproteobacteria*, *Alphaproteobacteria*, or *Verrucomicrobia*, corresponding to Type I, II, and III methanotrophs, respectively^12–13^. Previous research into the different growth conditions of Type I versus Type II methanotrophs found that Type II methanotrophs generally dominate high CH_4_, low oxygen environments along with nitrogen- and copper-limiting conditions^14–16^. However, Type II methanotrophs have also been found in soils with low CH_4_ concentrations^17–19^ likely due to two isoenzymes of the particulate methane monooxygenase that have different affinities for CH_4_^20^ making them more versatile metabolically.

Researchers have focused on the impact of rainforest-to-pasture conversion on CH_4_ cycling for decades^21–23^. Measurements of in-field gas flux generally show soil CH_4_ consumption across seasons in mature rainforest, while pasture soils emit CH_4_^24–25^. Over the last decade, further research into how tropical land use change influences CH_4_ cycling microorganisms found varied results. One study observed that the functional biomarkers for methanotrophy (*pmoA* and *mmoX*) decreased in cattle pastures with no change to the methanogenesis biomarker (*mcrA*), while another study observed a decrease in *pmoA* abundance from Type II methanotrophs and an increase in *mcrA* in cattle pastures^6,19^. These previous studies investigated how land use change in the Brazilian Amazon impacts the genomic potential of the soil methane-cycling microbial community, but no study has directly targeted the active community.

Metatranscriptomics, metaproteomics, and stable isotope probing are increasingly common techniques to target the active microorganisms in an environmental sample^26–29^. Previous research by our group using metatranscriptomics and metaproteomics was unable to determine if soil CH_4_ cycling genes and proteins were differentially expressed between land use types due to low counts (unpublished data). Therefore, for this study we used stable isotope probing to determine the active fraction of the soil microbial community cycling CH_4_, referred to henceforth as members of the active community. Stable isotope probing is commonly applied to study CH_4_ cycling in soil given the specific nature of the substrate and its relevance to climate change^30–32^. This technique uses the less abundant isotope of an atom, such as ^13^C-carbon, to label the microorganisms capable of consuming the ^13^C and, via their anabolic metabolism, incorporating it into their DNA, which then can be separated by ultracentrifugation from the community DNA. Subsequently, next generation sequencing enables the identification of active community members and provides insight into their functional potential.

The central goal of this study was to determine how the members of the active CH_4_-cycling microbial community, their functions, and CH_4_-related metabolic pathways changed across land use types (primary rainforest, cattle pasture, and secondary rainforest) and geographically distinct regions of the Brazilian Amazon. We hypothesized that the cause of increased soil methane production in cattle pastures was caused by a decrease in active methanotrophy. To test this hypothesis, we sampled sites at the most active deforestation frontiers in northeastern and southwestern Amazonia in the states of Pará (Tapajós National Forest) and Rondônia (Fazenda Nova Vida), respectively. To determine the community composition and functions of the active methane-cycling microorganisms, we coupled stable isotope probing (DNA-SIP) with metagenomics, using either ^13^C-labeled methane (CH_4_), carbon dioxide (CO_2_), or sodium acetate (NaAOc) as a substrate. Overall, we observed significant shifts in the active microbial community compositions and their methane-cycling functional genes between land-use types, geographic location, and substrates. Specifically, the abundance and diversity of active methanogens increased with conversion to pasture. Ultimately, this significant increase in active methanogens in pasture soils significantly correlated with the in-field methane gas flux. Therefore, we conclude that an increased abundance and diversity of active methanogens is causing the overall net positive methane flux in cattle pastures.

## METHODS

### Site description and sampling

At each sampling site, we first took soil gas flux measurements as detailed below, followed by collecting 5 to 6 soil cores. Seventy-two soil cores (5 cm diameter × 10 cm depth) were collected from the Tapajós National Forest and its adjacent areas in the State of Pará in June 2016 for DNA-SIP. Another 72 soil cores were collected from Fazenda Nova Vida and its adjacent areas in the State of Rondônia in April 2017 for DNA-SIP (GPS coordinates in Supplemental Methods). For each location, 18 soil cores were collected from each land use type, two primary rainforests (PF1 or PF2), one cattle pasture (P), and one secondary rainforest (SF). Soil cores were collected along a transect ranging from 100 to 200 meters with five equidistant sampling points. Three adjacent soil cores were taken from each sampling point with a fourth soil core taken from sampling points 2, 3, and 4 along the transect (Supplemental Figure 1). The cores were stored at 4°C until incubation with stable isotopes in the laboratory (up to 2 weeks later due to travel). The two additional soil cores collected at each sampling point were immediately homogenized, and then divided into two volumes for total prokaryotic community analyses from extracted DNA and for soil physical-chemical characterization.

### Stable isotope probing

During incubation with stable isotopes, the intact soil cores (∼200 g depending on soil density) were stored in gas-tight glass jars in the dark. For each combination of location (Pará or Rondônia) and transects across land use type (two primary rainforests, one pasture, one secondary rainforest), five soil cores were incubated with ^13^C-substrate and one additional core with ^12^C-substrate as the control. For each sampling site this resulted in a total of six cores for each of the three substrates, or 72 cores total for each location tested. Soils were incubated at 25°C for ∼7 months due to the low gas exchange at the surface top of the undisturbed soil column (20 cm^2^) compared to homogenized soil (20-32x lower rates; unpublished data). Either 25 mL of ^13^C-carbon dioxide (3% headspace concentration), 1 mL of ^13^C-sodium acetate (1 mM final concentration, added to the top of each soil core), or 25 mL of ^13^C-methane (3% headspace concentration) were added every two weeks. Equal volumes (1 mL) of sterile water were added to carbon dioxide and methane incubations. Air was added once a week to the ^13^C-methane incubations to ensure an oxic headspace. Pressure was released periodically prior to substrate injection from all jars. The duration of incubation was determined by monitoring the methane gas flux and attempting to ensure 20 mM of substrate was incorporated, following published recommendations to apply 5-500 µM ^13^C per g of soil^33^. Our target was to incorporate ∼100 µM ^13^C per g of soil, rendering shorter incubation times insufficient. Methane production or consumption was monitored throughout the incubation experiment by gas chromatography (Shimadzu GC-17A, Kyoto, Japan). After incubation, each soil core was sectioned longitudinally into five 2-cm tall segments (numbered 1-5 from top to bottom) and stored frozen at −20°C until DNA extraction.

### DNA extraction, quantification, and sample processing

DNA was extracted from 0.25 g of soil from all segments from two of the five ^13^C soil cores using the DNeasy PowerSoil DNA Extraction kit (Qiagen, Hilden, Germany) to determine the segment with the highest abundance of methanogens or methanotrophs based on the respective functional marker genes using qPCR as described below. Upon identifying the segment with the highest genomic abundance of methanogens or methanotrophs, DNA was extracted from 4 g of soil from the identified segment of three ^13^C soil cores and from the ^12^C-control for each substrate/sample site combination using the DNeasy PowerMax Soil Kit (Qiagen). DNA was quantified fluorometrically using the Qubit dsDNA Broad-Range assay (Invitrogen, Carlsbad, CA). A total of 5 µg of DNA was subjected to ultracentrifugation according to a previously described protocol^33^, followed by fractionation of the density gradient into 12 fractions of equal volume. The continuity of the density gradient was confirmed with a refractometer. DNA was precipitated following the published protocol^33^ except for the addition of 20 µg linear acrylamide (Invitrogen) instead of glycogen and each fraction was quantified using fluorometry via the Qubit dsDNA High-Sensitivity assay (Invitrogen). To identify the fractions with ^13^C-labeled DNA, we quantified the abundance of methanogens or methanotrophs in each fraction using qPCR of the respective functional gene marker for a subset of samples compared to their respective ^12^C-controls (details in Supplementary Methods). We pooled the ^12^C (∼1-5) and ^13^C (∼6-12) fractions, respectively. Since the GC content of microbial DNA can influence DNA density, we sequenced both the light and heavy DNA fractions from our ^12^C-controls for a total of 12 ^12^C-light DNA, 12 ^12^C-heavy DNA, and 36 ^13^C-heavy DNA samples per location.

### Quantitative PCR

The particulate methane monooxygenase alpha subunit gene (*pmoA*) was amplified using the primer pair A189f/mb661r^34–35^, and the gene for the methyl coenzyme M reductase alpha subunit (*mcrA*) was amplified using the primer pair mlas/mcra-rev^36^. Standard reaction mixtures and thermocycler conditions are specified in Supplemental Methods.

### Sequencing

All DNA library preparation and sequencing were performed at the University of Oregon Genomics and Cell Characterization Core Facility (Eugene, OR) (Supplementary Methods). Briefly, the three genes of interest (16S rRNA, *pmoA*, and *mcrA*) were amplified using custom dual-indexed PCR primers designed by the core facility. For each location, paired-end 300 bp amplicon sequencing of the pooled heavy fractions for three ^13^C-samples per sample site/substrate combination and the pooled heavy and light fractions for all ^12^C-controls was completed on an Illumina MiSeq sequencer (Illumina, San Diego, CA). For metagenomes, sequencing of the heavy fraction of two ^13^C-samples per sampling site/substrate combination and all ^12^C-controls was performed on an Illumina HiSeq4000 across two flow lanes for each location. All sequences were demultiplexed at the core facility.

### Soil physical-chemical analysis

Homogenized soil samples stored at 4°C were processed as described in detail previously^37^.

### Methane Gas Flux Measurements

In-field soil CH_4_ fluxes were measured using a field-deployable Fourier transform infrared spectrometer (Gasmet, DX 4015, Vantaa, Finland) sampling a recirculating flow-through soil flux chamber placed on soil collars (aluminum, inner area of 284 cm^2^) that were installed in soil (5 cm deep) at least 20 minutes beforehand. Fluxes were calculated from the rate of headspace CH_4_ accumulation or depletion in a 30-minute period. We fit linear models to concentration vs. time trends, limiting the extend of the time series used to the initial linear decline in cases when nonlinear behavior was observed at late stages in the measurement^38–39^.

### Data and statistical analysis

Amplicon sequences were processed and analyzed using the DADA2 pipeline in QIIME2^40–41^. Metagenome sequences were processed and annotated using MG-RAST^42^. GenBank and SEED Subsystem were used for the organismal and functional annotations, respectively. The SEED Subsystem annotation “Methanogenesis strays” is described as “several additional genes and clusters from methanogens.” The influence of homogeneous dispersion within each sample and heterogeneous dispersion between samples was assessed using the Permdisp and Adonis functions, respectively, from the ‘vegan’ package in R on dissimilarity matrices made from the annotation tables^43–44^. To specifically target the active microbial community, the metagenomic annotations were rarefied (vegan:rrarefy) and counts were normalized to the ^12^C-control for each substrate (see Supplementary Methods). After rarefication, the dissimilarity between land use types within each substrate were analyzed (vegan:Adonis). STAMP v2.1.3 was then used to identify active microorganisms and functions by comparing each individual ^13^C-sample to their respective ^12^C-control using Fisher’s exact test^45^. All figures were made in R v3.5.1 using ggplot2^43,46^. Soil physical-chemical data were analyzed using ANOVA with a Tukey-Kramer post-hoc test. Correlation analyses between the in-field methane flux and the abundance of active methane-cycling taxa or functional annotations were completed using a Pearson correlation (cor.test)^43^.

#### Active Fraction Analysis

In this study, “active” means that the cells were actively growing (anabolically incorporating ^13^C) and not just metabolically active (catabolic turnover of ^13^C substrate independent of growing). The incubations with ^13^C-labeled substrates determine both actively growing and metabolically active community members, and we used our ^12^C incubation controls to correct for the metabolically active part. Therefore, an annotation was deemed active, if it was significantly higher (p < 0.05) in the ^13^C-sample compared to the ^12^C-control. Samples were normalized to their respective ^12^C-control with features that had less abundance in the ^13^C- than ^12^C-samples being marked as 0 counts. Samples from the same substrate were compared between land use types in STAMP using the multigroup stats function (ANOVA with Tukey-Kramer Post-hoc test). All significantly different annotations were checked again to see if they were active in the samples.

### Data Accessibility

Metagenomes are available publicly on MG-RAST under the following project accession numbers: mgp88468 and mgp86794. All raw amplicon sequence files have been deposited on figshare under the following DOI: 10.6084/m9.figshare.10565552, 10.6084/m9.figshare.10565690, 10.6084/m9.figshare.10565870, 10.6084/m9.figshare.10565897, 10.6084/m9.figshare.10565957, 10.6084/m9.figshare.10565801.

## RESULTS AND DISCUSSION

### Active Methane-Cycling Community Changes with Land-Use

To understand the active methane-cycling microbial community composition and abundance, we analyzed sequences of both PCR-amplified marker genes (16S rRNA, *mcrA*, and *pmoA*) and metagenomes. The amplification-based approach makes our data comparable to many microbial studies that use these biomarkers, but this method comes with the potential issues of primer bias allowing for missed taxonomic groups, lower phylogenetic resolution, and no additional information on ecosystem processes^47–48^. Therefore, after confirming that enough label was present in the target sample using amplicon-based sequencing, we used metagenomics to gain a deeper understanding of the ^13^C-labeled methane-cycling community and its supporting members^49–51^. The composition of the total soil microbial community, based on 16S rRNA, significantly differed between geographic locations (Rondônia vs Pará; p=1e-03 r^2^=0.118), individual land-use types (primary rainforest(PF), pasture(P), secondary rainforest(SF), p=1e-03 r^2^=0.08), and added substrates (CH_4_, CO_2_, NaAOc, p=1e-03 r^2^=0.08) (Figure 1). When we specifically targeted the active community, we found that only samples incubated with CO_2_ significantly differed between locations (Pará CO_2_ p=3e-03; Rondônia CO_2_ p=1.8e-02; Supplemental Figures 2-3). It was unsurprising that location is the main driver to differentiate the total microbial community since Rondônia and Pará are separated by ∼1500 km. Also, abiotic factors such as seasonal differences and/or soil physical-chemical properties could be driving these locational differences^52–54^. The significant differences in CO_2_ incubated samples may be due to the similarity of the overall active community across samples, while the methane-consuming or -producing community makes up only a small fraction of that community and the signal is lost when we look at the community composition broadly.

**Figure 1.**
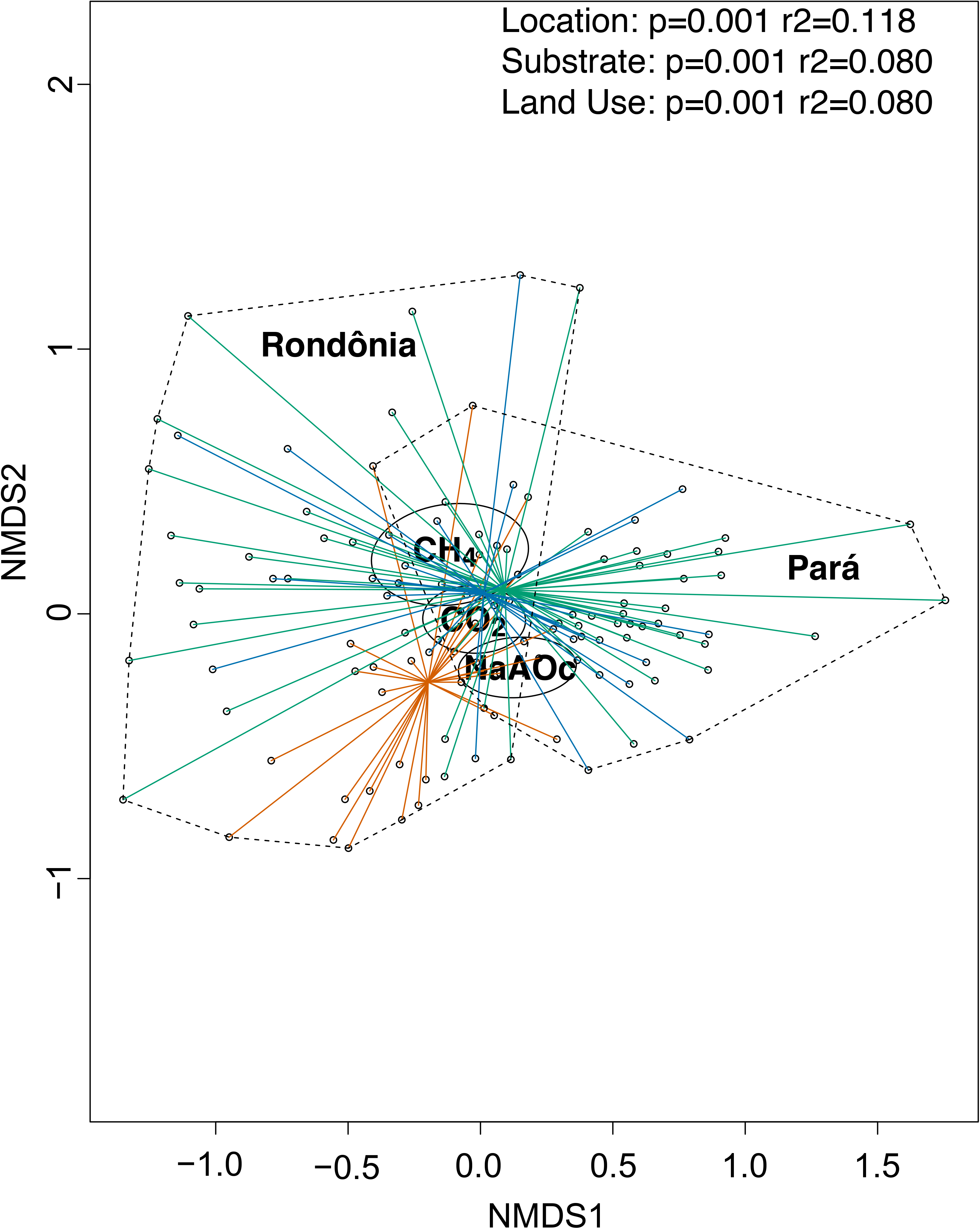
Composition of the total soil microbial community based on 16S rRNA. Non-metric dimensional scaling plot of rarefied 16S rRNA SILVA annotations at genus-level from all samples. The dotted lines outline samples from each geographic location (Pará and Rondônia). The colored lines connect samples from the same land-use type to the centroid (primary rainforest = green, pasture = orange, and secondary rainforest = blue). The circles represent the standard error dispersion of samples for each substrate (NaAOc=sodium acetate, CO_2_=carbon dioxide, CH_4_=methane). The p-values and r^2^ values for each variable (Location, Substrate, Land Use) are derived from the Adonis function in the vegan package.

When we investigated the richness of active methane-cycling communities, we found that pasture samples had the highest active methanogen richness in metagenomes from both locations and regardless of substrate (CO_2_ or NaAOc); however, it was only significant in Rondônia NaAOc samples (P vs PF p=9.6e-03, P vs SF p=7.9e-03; Figure 2, Supplemental Table 1). All active methanogens that significantly changed abundance between land-use types were associated with pasture soils in both locations (Table 1). Specifically, *Methanosarcina* spp. dominated the active methanogens for most samples in both locations regardless of substrate (Figure 3, Supplemental Tables 2-3; 5-6). These archaeal species are known to have multiple methanogenesis pathways making them capable of utilizing both ^13^CO_2_ and ^13^NaAOc, likely explaining their dominance in both locations and substrates^55–58^.

**Figure 2.**
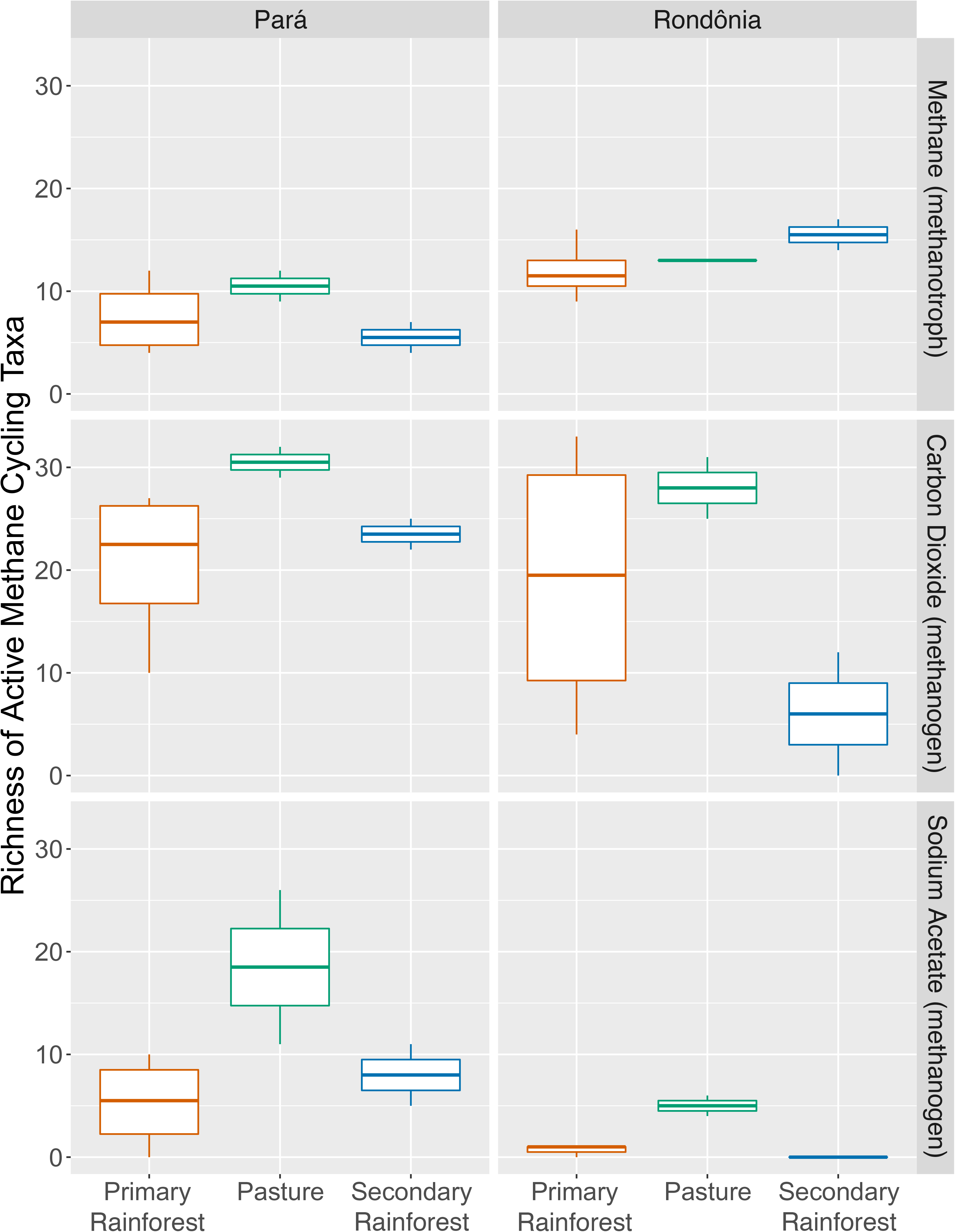
Richness of active methane cycling taxa (methanotroph or methanogen) from two geographic locations (Pará or Rondônia), three land-use types (primary rainforest = green, pasture = orange, and secondary rainforest = blue) incubated with one of three substrates (methane, carbon dioxide, and sodium acetate).

**Figure 3.**
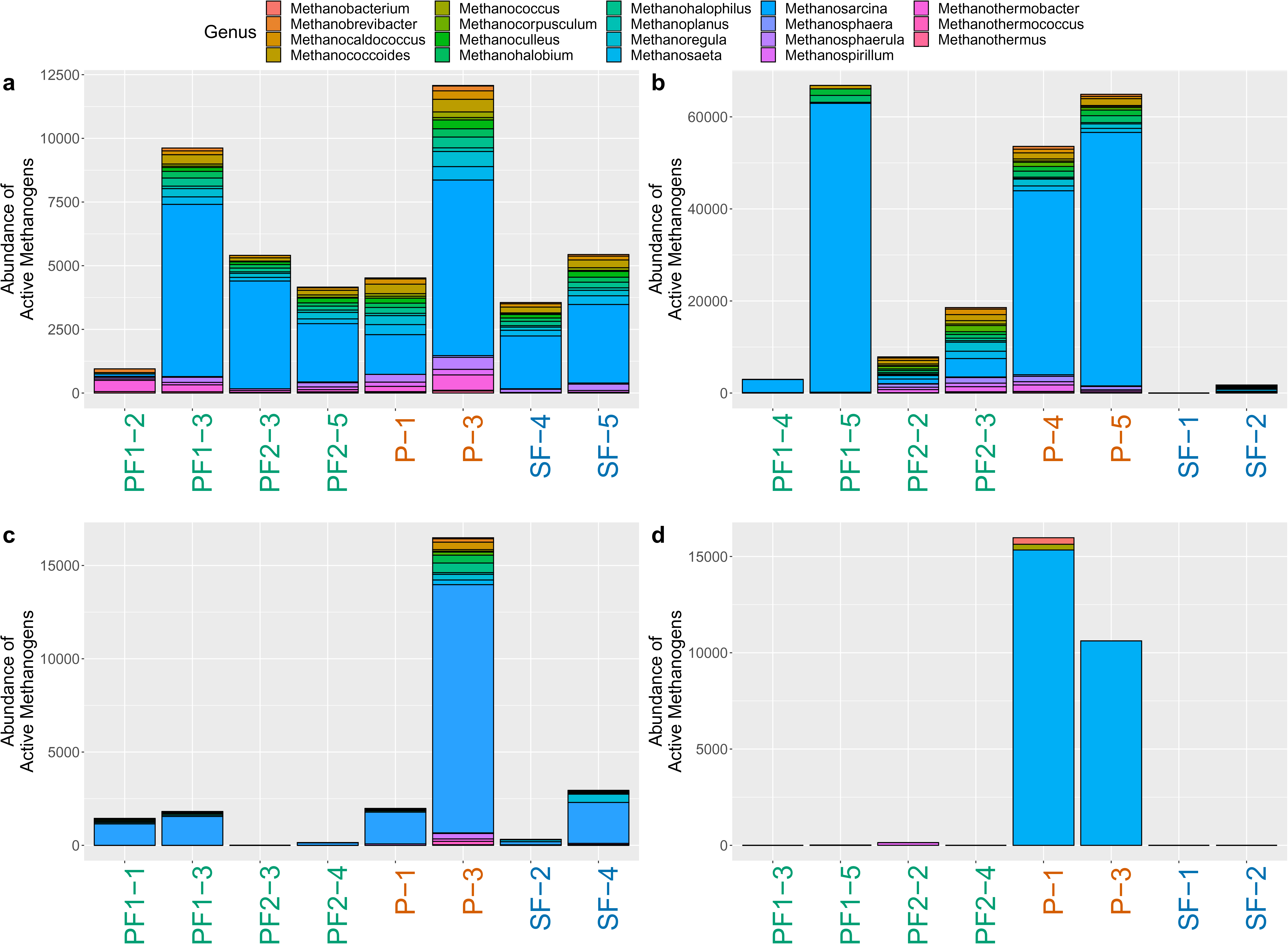
Abundance of active methanogen genera in the ^13^C metagenome samples. (a) The abundance of active methanogens in soils from Pará incubated with ^13^CO_2_, (b) the abundance of active methanogens in soils from Rondônia incubated with ^13^CO_2_, (c) the abundance of active methanogens in soils from Pará incubated with ^13^NaAOc, (d) the abundance of active methanogens in soils from Rondônia incubated with ^13^NaAOc. Samples on the x-axis are colored by land use type (primary rainforest (PF) = green, pasture (P) = orange, and secondary rainforest (SF) = blue).

**Table 1.** Methanogens and methanotrophs that are both active and significantly different between land use types (primary rainforest, pasture, secondary rainforest). The term ‘Land use association’ signifies which land use is associated with a significantly higher abundance of the taxon. Mean relative abundance (%) depicts the average relative percent of each taxon in each land use.

We observed a significantly higher abundance of total active methanogens in Rondônia pasture soils compared to both primary and secondary rainforest samples in ^13^NaAOc samples (p=1e-03, p=3.8e-02, respectively) and compared to secondary rainforest in ^13^CO_2_ samples (p=9e-03). There was no significant difference in the abundance of total active methanogens between land-use types for either methanogenic substrate in Pará, but many taxa did significantly change abundance (Table 1). Previous research studies showed mixed findings on methanogen communities response to tropical land-use change ranging from no change to increased *mcrA* gene abundance in pastures^59,6,19^. By targeting the active community, we directly show that pasture soils have a higher richness of active methanogens and specific methanogenic taxa significantly increase abundance. This increase in methanogen abundance and richness is likely due to the increased soil carbon cycling occurring in pasture soils^60–61^.

Previous research into methanotrophy across Amazonian land uses found methanotroph abundance to be lower in pasture relative to primary forest soils^6,59,19^. Based on these studies, we hypothesized that pasture soils would have the lowest abundance and richness of active methanotrophs. Unlike the active methanogen community, we did not find a consistent association between active methanotroph richness and land-use types across locations. The highest richness was either found in pasture or secondary rainforest for Pará and Rondônia, respectively, but it was not significant (Figure 2). The total active methanotroph abundance did not significantly change between land-use types. In both locations and all land-use types, Type II methanotrophs (Alphaproteobacteria) dominated the active methanotroph community (Figure 4, Supplemental Tables 4, 7). Only one primary rainforest sample from Pará was dominated by Type I methanotrophs and only one Type III methanotroph was found to be active, *Methylacidiphilum*, but remained rare (Supplemental Tables 4, 7).

**Figure 4.**
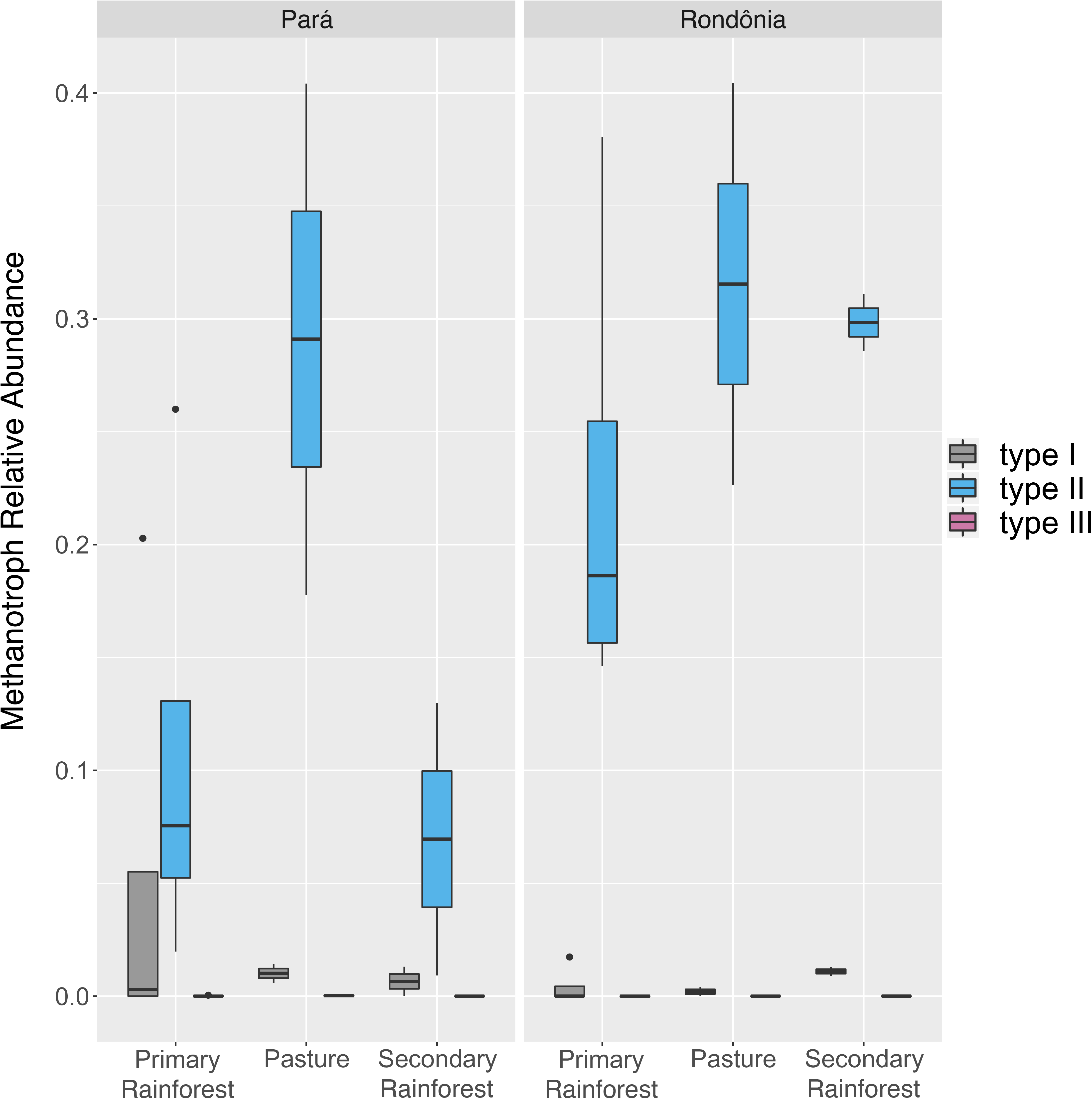
Relative Abundance of methanotroph types I, II, and III across both geographic locations (Pará and Rondônia) and land use types (primary rainforest, pasture, and secondary rainforest). Type I = grey, Type II = blue, Type III = pink.

Although, the total abundance of active methanotrophs did not significantly change between land-use types, the abundance of specific methanotrophs changed in Pará and Rondônia associating with pasture and secondary rainforest, respectively (Table 1). This was surprising and not what we hypothesized based on previous studies^6,19,59^. Several factors should be considered to address this discrepancy. First, our study targeted the microorganisms actively consuming CH_4_ rather than looking at the total microbial community. Studies of the total microbial community can be influenced by the potential presence of extracellular DNA, which may affect estimates of abundance and diversity^62–64^. Additionally, we incubated our samples at CH_4_ concentrations above those in the atmosphere due to the inability to label the community at low concentrations. Although necessary for the technique, this could influence the composition and activity of the CH_4_-consuming community. Furthermore, there is a possibility that we incorrectly assumed primary rainforests would have the highest methanotroph richness and abundance since these forests are known to be methane sinks^21–22^. Here, our findings enable the next question to test what environmental variables govern the active CH_4_-consuming community between land-use types in the Brazilian Amazon.

### Dominant Active Methanogenesis Pathways Differed Between Locations

We next asked which CH_4_-related metabolic pathways were active across land-use types and how they changed in response to deforestation. We observed active methanotrophy based on the abundance of the genes for particulate methane monooxygenase (*pmmo*) and soluble methane monooxygenase (*smmo*) in all ^13^C-labeled samples and in both locations (Figure 5; Supplemental Table 8). The *pmmo* genes were abundant and active in 94% of samples while *smmo* was active in most secondary rainforest samples and Rondônia-PF1 (Figure 5; Supplemental Table 8). The secondary rainforest and pasture samples at Rondônia significantly increased in *pmmo* abundance (p=6e-03; p=3e-02, respectively) compared to primary rainforest (Supplemental Table 8). We found no significant difference in the abundance of any active methanotrophy-related genes across land-use types in Pará (Figure 5). Overall, Rondônia had a significantly higher relative abundance of *pmmo* to total methanotrophy annotations compared to Pará (Supplemental Table 8) (p=1e-04). Soil physical-chemical properties are known to influence the activity of these different methane monooxygenases^65^. Copper is a key component regulating the activity and abundance of these methane monooxygenases having a positive relationship with *pmmo* abundance ^66–67,12^. We observed a significantly higher concentration of copper (9x) in Rondônia compared to Pará (p=7.42E-06) which may explain the increased abundance of active *pmmo* genes.

**Figure 5.**
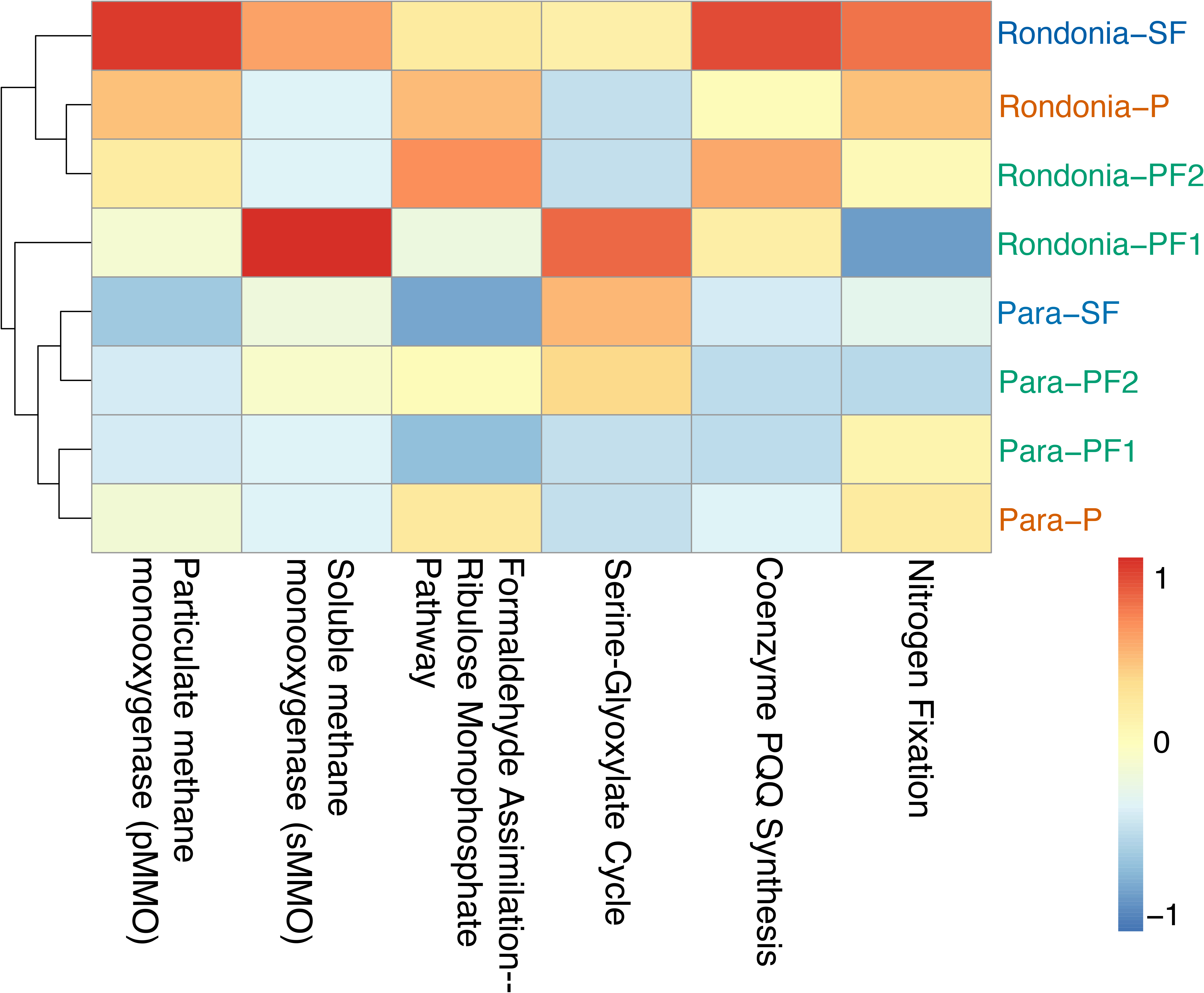
Heatmap visualizing the average relative abundance of active genes involved in methanotrophy pathways. The scale is from lowest relative abundance (blue) to highest relative abundance (red) of the genes and is normalized to each gene (i.e. column). The metagenome samples are on the y-axis are colored by land use type (primary rainforest (PF) = green, pasture (P) = orange, and secondary rainforest (SF) = blue) and have the location (Rondônia or Pará) in the label. The genes involved in methanotrophy are on the x-axis. The dendrogram shows the Euclidean distance between samples.

Regardless of location, the abundance of active methanogenesis genes dominated in pasture compared to other land-use types. Interestingly, in Pará we observed these significant increases in the ^13^CO_2_ incubation, while in Rondônia the ^13^NaAOc incubation accounted for the increased abundance (Figure 6; Supplemental Table 9). The Pará ^13^NaAOc incubation presented some significant changes in methanogenesis-related genes including Coenzyme F420 synthesis (p=8e-05), methanopterin biosynthesis2 (p=2e-03), and methanogenesis strays (p=1e-03). The two pasture samples in Pará ^13^NaAOc incubations performed very differently. Although pastures are considered to be more biotically homogeneous^68^, these two samples differed strongly with one sample having about 8.5x more active methanogens (Supplemental Table 4). When the relative abundance of active methanogenesis genes to total annotations was investigated, we identified a significant difference between land-use types in the Pará ^13^CO_2_ incubations (PF v P p=1e-03, SF v P p=1.8e-02) and in the Rondônia ^13^NaAOc incubations (PF v P p=7e-02, SF v P p=1.8e-02) (Supplemental Table 10). In addition to methanogenesis genes changing between land-use types, we observed an increase in carbon cycling activity in Pará pasture soils incubated with ^13^CO_2_ (Glycolysis and gluconeogenesis p=1e-03, Pentose phosphate pathway p=2e-03, Entner-Doudoroff pathway p=3e-02). Overall, we found that active methanogenesis was driven by methanogens using the hydrogenotrophic pathway in Pará and the acetoclastic pathway in Rondônia (Supplemental Table 9). This shift in the dominant methanogenesis pathway between locations may be due to differences in the physical-chemical soil parameters or a result of the types of fermentation leading to either more acetate or hydrogen production. Interestingly, the active methanogen community was dominated by *Methanosarcina* spp. in both locations. Members of the genus *Methanosarcina* are known to require three different types of hydrogenases for the reduction of CO_2_ to CH_4_ with electrons derived from H_2_^58^. The significantly increased activity of multiple types of hydrogenases (Energy conserving hydrogenase ferrodixin Ech p=1.6E-08; membrane bound hydrogenases p=4.6e-02; Archaeal membrane bound hydrogenases p=0.048; Coenzyme F420 hydrogenase p=5e-02) in soils from Pará compared to soils from Rondônia indicates a selection for the hydrogenotrophic pathway. This selection is supported by the increased availability of trace metals (iron) in soils from Pará which are needed by methanogenic hydrogenases^58^.

**Figure 6.**
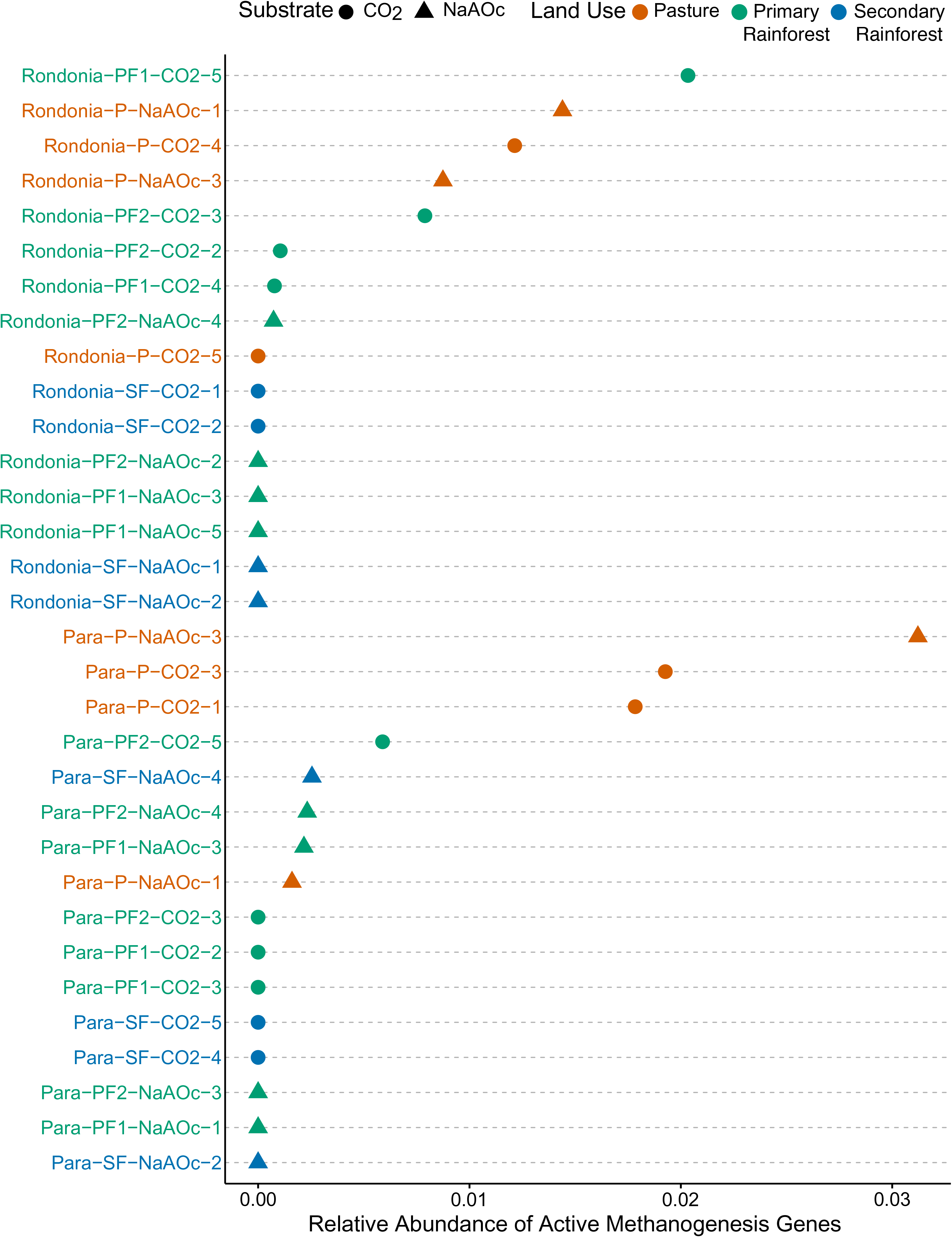
Dot chart illustrating the relative abundance of active methanogenesis genes. The samples are grouped by location in descending order and include both methanogenic substrates (CO_2_ and NaAOc). The colors correspond to the land use type (green = primary rainforest, orange = pasture, blue = secondary rainforest). The shapes of the dots correspond to substrate (circle=CO_2_, triangle = NaAOc). Active methanogenesis genes includes SEED subsystem annotations as “Methanogenesis,” “Methanogenesis from methylated compounds,” and “Methanogenesis strays,” Methanogenesis strays are “additional genes and clusters from methanogens”. The specific genes associated with “Methanogenesis strays” can be found by searching for the subsystem on the SEED viewer (http://rast.theseed.org/FIG/seedviewer.cgi?page=SubsystemSelect).

### Land-Use Change Alters Key Redox-Cycling Active Taxa

In the context of highly complex soil microbial communities, methanogens need other microorganisms to produce the substrates necessary for this redox reaction to occur. Methanogenesis is one of the least thermodynamically favorable anaerobic reactions; therefore, other redox reactions must transpire prior to methanogenesis^69^. Although we were targeting active methane-cycling microorganisms in this study, the methanogenic substrates used, ^13^CO_2_ and ^13^NaAOc, are not exclusively used by methanogens. Therefore, we investigated which coexisting microorganisms were actively consuming these substrates and thereby interacting with methanogens. Many non-methanogenic but active microbial taxa changed significantly in abundance between land use types in both geographic locations. In the Pará ^13^CO_2_ SIP incubations, we observed a significant increase of active *Syntrophus aciditrophicus* (p=1E-06) in pasture along with many known sulfate-reducing bacteria (Supplemental Table 10. *Syntrophus aciditrophicus* is known to promote the growth of *Methanospirillum* spp., which accounted for 2.76% of the active methanogen community in Pará ^13^CO_2_ pasture samples^70^. In the secondary rainforest, we found a significantly higher abundance of various active nitrifying and sulfur-oxidizing bacteria, such as *Nitrobacter* and *Thioalkalivibrio*. Many of these microbial groups are known to utilize CO_2_ and have thermodynamically preferred redox potentials^71–76^. Rondônia pastures increased in active ammonia-oxidizing microorganisms including *Nitrosococcus* and *Geobacillus* species (^13^CO_2_ samples; Supplemental Table 10). One potential cause of increased *Geobacillus* species is the slash and burn process used to create pastures that deposits hydrocarbons in the soil, which these microorganisms are known to use^77–79^.

The abundance of active *Geobacillus*, *Clostridium,* and *Sulfolobus* spp. increased in Pará ^13^NaAOc incubated pasture soils (Supplemental Table 11). Some *Geobacillus* and *Clostridium* spp. are known to utilize acetate, which may explain their increased abundance in the ^13^C-labeled community^80–81^. The denitrifying bacterium *Hyphomicrobium denitrificans* was active and significantly increased abundance in Pará primary rainforest samples along with the genes associated with denitrification (p=0.02). In the ^13^NaAOc Rondônia soils, we observed a significant increase in both sulfate-reducing and sulfur-oxidizing microorganisms along with nitrate-reducers in secondary rainforest with many competitors for acetate as a carbon source^82,74^ (Supplemental Table 11). It is well documented that before methanogenesis is able to occur nitrate and sulfate must be depleted as electron acceptors^69^. The increased abundance of active sulfate and nitrate reducers in the Rondônia secondary rainforest and overall lack of active methanogenesis indicates that these more favorable electron acceptors were still available in the soil during incubation with ^13^NaAOc inhibiting methanogenesis through substrate competition^83–84^.

### Soil Physical-Chemical Parameters Increase Potential Methane Production

Land-use change is one of the strongest drivers to alter soil ecosystems. Parallel changes to the soil physical-chemical parameters, physical structure, and aboveground vegetation may provide additional support for increased methanogenesis in pasture soils. Specifically, the compaction caused by cattle grazing creates more anoxic microsites providing more opportunity for methanogenesis to occur^85^. The comparison of soil physical-chemical parameters between the geographic locations presented several significant differences (Supplemental Tables 12-13). Of note were increased concentrations of sulfur (p=2.95E-15) and copper (p=7.42E-06) along with higher pH (p=1.35E-07) in Rondônia compared to Pará, and total soil acidity (p=9.29E-11) and total nitrogen (p=2.31E-06) were significantly higher in Pará soils. For both locations, the soil pH was significantly higher in pasture compared to primary rainforests. Soil bulk density was found to be highest in pasture from both locations (Supplemental Figure 5). The increased pH in pasture soils likely helps support methanogenesis since optimum process activity is at near neutral pH and quickly decreases as the pH becomes more acidic^86^. Another contributing factor to the increased methanogenesis in pasture soils is due to *Urochloa brizantha* excreting large amounts of carbon as root exudates into the soil^87^. With increased carbon availability in pasture soils, there is overall increased soil microbial activity^88^. All of these changes to the soil in pastures could contribute to the increased methanogenic activity observed in our SIP study.

### Relating Functional Activity from In-Field Methane Gas Flux to Functional Potential from Metagenomics

Research into modeling biological activity to help predict future climate scenarios has continued to grow over the past few decades^89–91^. Since this study focused on the active methane-cycling microbial community across land use types, we performed a correlation between the abundance of the active community or of functional biomarkers with the gas fluxes measured in our sites. Tentatively, we found the strongest correlation between total active methanogen abundance and field methane flux (p=6e-04, cor=0.573) (Figure 7). However, these findings will need to be further investigated across more sampling points to confirm their validity broadly. Overall, only Rondônia showed significant differences in field methane flux across land-use types, with pasture having significantly higher methane emissions (PF vs P p=1e-04, SF vs P p=2e-03, SF vs PF p=9.9e-01; Supplemental Figure 6). The correlation observed between active methanogen abundance and methane flux in the field provides further support that an increase in methanogenesis activity is driving the change in methane flux between land-use types.

**Figure 7.**
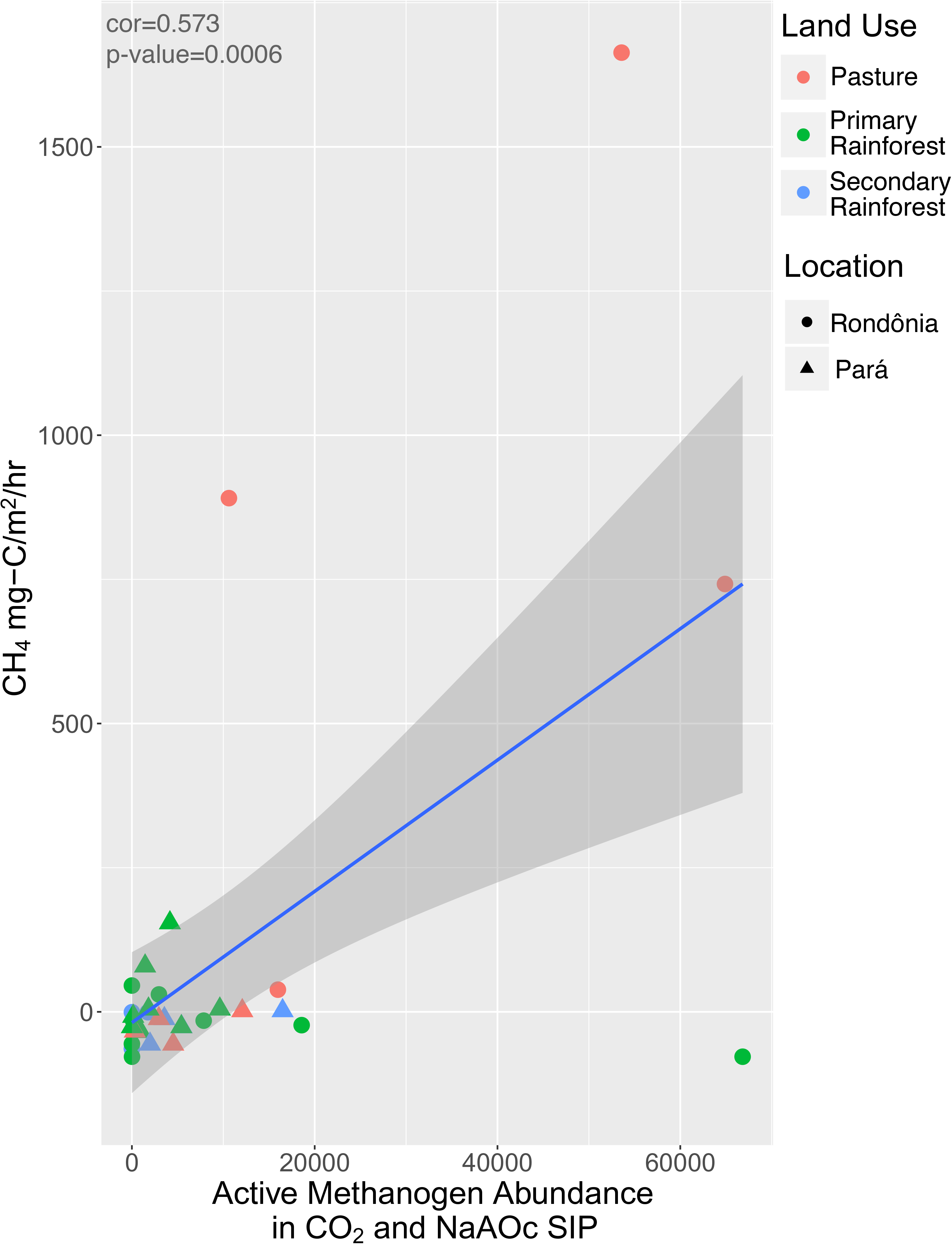
The field gas flux for methane (CH_4_ mg-C/m^2^/hr) correlated to active methanogen abundance in ^13^CO_2_ and ^13^NaAOc SIP incubations. Primary rainforest = green, pasture = red, and secondary rainforest = blue. Samples from Rondônia are circles and samples from Pará are triangles.

### Minimal Enrichment of Methane-Cyclers After Incubation

In SIP studies specifically, we target the active microbial groups involved in a biogeochemical process; however, the majority of soil SIP studies use homogenized soil where soil columns get sieved^92–97^. It is clear from the literature that soil structure is an important aspect of microbial activity and carbon cycling; therefore, if possible, microbial activity should be studied under environmentally-relevant conditions. This study shows the feasibility of keeping soil and its assembled microbial communities more similar to the natural environment by incubating soil cores in-tact. We observed that even after seven months of incubation, the abundance of functional marker genes (*pmoA* and *mcrA*) did not become greatly enriched. Compared to field soils at the time of sampling, there was a small but significant increase of *mcrA* gene copies in Pará ^13^CO_2_ SIP soil (p=4.6e-03), but no significant difference in soils from Para amended with ^13^NaAOc (p=7.2e-01) (Supplemental Figure 7). Overall, there was no significant difference between the Rondônia SIP and field soils’ *mcrA* gene abundance. The only significant difference found was between ^13^CO_2_-incubated primary forest and pasture samples (p=4.6e-02). Interestingly, *pmoA* gene abundance decreased significantly in SIP incubated soils from Pará (p=1E-05) and Rondônia (p=4E-07, Supplemental Figure 8). One possible explanation for the decreased *pmoA* gene abundance between SIP incubated and field soil is that during the incubation the methanotrophic community was potentially altered. Our comparative analysis of the metagenome data supports this possibility as a 7.7-fold and 4.0-fold increase from Rondônia and Pará, respectively, were observed in obligate methanotroph abundance between ^13^C vs ^12^C heavy fraction samples. Since primer bias is a common problem, as previously discussed, the change in community could alter the compatibility of the primer to the *pmoA* sequences of the changed community; thus, potentially presenting a lower *pmoA* abundance in the SIP than field soils.

## CONCLUSIONS

Land use change from rainforest to pasture stimulates the soil methanogenic community in the Brazilian Amazon. Using undisturbed soil columns for SIP incubations, we were able to ascertain that methanogen abundance and activity is significantly higher compared to both primary and secondary rainforests which could drive methane emissions from the soil of Brazilian cattle pastures. Future studies should focus on identifying what specific environmental factors are responsible for increased methanogenesis in pasture soils (i.e. pH, vegetation, compaction, carbon or trace element availability, etc.), so that land management can better mitigate CH_4_ emissions. Another important finding was that secondary rainforests in both locations have recovered as CH_4_ sinks with an active methanotrophic community. Through large forest restoration efforts occurring in the tropics, there is potential to see these forests recover with enough time to overcome excess CH_4_ production. It is currently unknown how long secondary rainforests take to recover as a CH_4_ sink, and how widespread this recovery is geographically. Adoption of best management practices in pastures can compensate for a small fraction of the impact of deforestation on net emission of greenhouse gases and the loss of carbon from Amazonia. With the currently accelerating expansion of land-use change in Amazonia understanding which players might assist mitigation of concomitant greenhouse gas production is increasingly important for all agricultural management.

## ACKNOWLEDGEMENTS

The authors thank the owners and staff of Agropecuaria Nova Vida for logistical support and permission to work on their property. We also thank all collaborating private landowners of Santarem for their support and access to their land. We would like to thank the Large-Scale Biosphere-Atmosphere Program (LBA), coordinated by the National Institute for Amazon Researchers (INPA), for the use and availability of data for logistical support and infrastructure during field activities. We are grateful to Erika Berenguer, Liana Chesini, and Jos Barlow, members of the EcoFor Project at Lancaster University, UK, for sharing some northern Amazon sites^98^ with us and for logistical support during field activities. Additionally, we are grateful to Alexandre Pedrinho and Wagner Piccinini for assistance with fieldwork and Kiran Khan, Alex Thompson, Luke Gibney, and Rachel Feldman for assistance processing samples.

## Supplemental Table and Figure Legends

Suppl. Figure 1. Visualization of the sampling scheme for each geographic location and land-use type (two primary rainforests, one cattle pasture, and one secondary rainforest). Soil cores (18 x) were collected along a transect ranging from 100 to 200 m with five equidistant sampling points. Three adjacent soil cores were taken from each sampling point with a fourth soil core taken from sampling points 2, 3, and 4 along the transect.

Suppl. Figure 2. Non-metric multidimensional scaling indicating how similar the active soil microbial communities are for the substrates (a) ^13^CH_4_, (b) ^13^CO_2_, (c) ^13^NaAOc from Pará. The plot was created using a Bray–Curtis dissimilarity matrix among all samples. Primary rainforest = green, pasture = red, and secondary rainforest = blue.

Suppl. Figure 3. Non-metric multidimensional scaling indicating how similar the active soil microbial communities are for the substrates (a) ^13^CH_4_, (b) ^13^CO_2_, (c) ^13^NaAOc from Rondônia. The plot was created using a Bray–Curtis dissimilarity matrix among all samples. Primary rainforest = green, pasture = red, and secondary rainforest = blue.

Suppl. Figure 4. Abundance of active methanotroph genera in samples from (a) Pará or (b) Rondônia incubated with ^13^CH_4_. The active metagenomes are colored based on land-use type with primary rainforest = green, pasture = orange, and secondary rainforest = blue.

Suppl. Figure 5. The soil bulk density (g/cm^3^) for each land use (primary rainforest, pasture, secondary rainforest) and geographic location (Rondônia and Pará).

Suppl. Figure 6. The field gas flux for methane (CH_4_ mg-C/m^2^/hr) correlated significantly with active methanogen abundance in SIP incubations for either substrate ^13^CO_2_ and ^13^NaAOc. Primary rainforest = green, pasture = red, and secondary rainforest = blue. Samples from Rondônia are circles and samples from Pará are triangles.

Suppl. Figure 7. The log *mcrA* gene copies per ng of DNA for soils from three land-use types (primary rainforest, pasture, and secondary rainforest) in Rondônia and Pará incubated with ^13^CO_2_, ^13^NaAOc, or no incubation (field soil). ^13^CO_2_ samples = red, ^13^NaAOc samples = blue, unaltered samples = green.

Suppl. Figure 8. The log *pmoA* gene copies per ng of DNA for soil from three land-use types (primary rainforest, pasture, and secondary rainforest) in Rondônia and Pará incubated with ^13^CH_4_ or no incubation (field soil).

Suppl. Table 1. Significance values (p-values) from ANOVA with Tukey Honestly Significant Difference Test comparing the richness of active methane-cycling taxa.

Suppl. Table 2. The abundance of active methanogen species found in each ^13^CO_2_ SIP incubation from Rondônia.

Suppl. Table 3. The abundance of active methanogen species found in each ^13^NaAOc SIP incubation from Rondônia.

Suppl. Table 4. The abundance of active methanotroph species found in each ^13^CH_4_ SIP incubation from Rondônia. The Type column specifies whether that methanotroph species is Type I, II, or III.

Suppl. Table 5. The abundance of active methanogen species found in each ^13^CO_2_ SIP incubation from Pará.

Suppl. Table 6. The abundance of active methanogen species found in each ^13^NaAOc SIP incubation from Pará.

Suppl. Table 7. The abundance of active methanotroph species found in each ^13^CH_4_ SIP incubation from Pará. The Type column specifies whether that methanotroph species is Type I, II, or III.

Suppl. Table 8. The relative abundance of active methanotrophy or methanotrophy-related genes for samples incubated with ^13^CH_4_. Location indicates whether the sample is from Rondônia or Pará. Land use states whether the sample is from a primary rainforest, pasture, or secondary rainforest. The within and between location values show the p-value from a two-tailed t-test comparing land-use types.

Suppl. Table 9. The relative abundance (%) of active methanogenesis genes (methanogenesis + methanogenesis strays + methanogenesis from methylated compounds) to the total methanogenesis gene annotations for each sample incubated with either 13CO2 or 13NaAOc. Location indicates whether the sample is from Rondônia or Pará. Land use states whether the sample is from a primary rainforest, pasture, or secondary rainforest. SIP Incubation indicates whether the sample was incubated with 13CO2 or 13NaAOc. PF = primary rainforest, P = pasture, SF = secondary rainforest. Methanogenesis strays are described by SEED Subsystem as “additional genes and clusters from methanogens”. The specific genes associated with these SEED Subsystems can be found by searching for the subsystem on the SEED Viewer (http://rast.theseed.org/FIG/seedviewer.cgi?page=SubsystemSelect). The two-tailed t-test values are p-values with significant (p<0.05) highlighted in red.

Suppl. Table 10. Microbial species implicated in the sulfur, nitrogen, or carbon cycle that were active and significantly different between land use types in Rondônia or Pará ^13^CO_2_ -supported SIP samples. The term “Land use association” indicates the land use type that had (1) a significantly higher abundance than the other land use types, and (2) the microbial species was active in that land use type.

Suppl. Table 11. Microbial species implicated in the sulfur, nitrogen, or carbon cycle that were active and significantly different between land use types in Rondônia or Pará ^13^NaAOc-supported SIP samples. The term “Land use association” indicates the land use type that had (1) a significantly higher abundance than the other land use types, and (2) the microbial species was active in that land use type.

Suppl. Table 12. Soil physico-chemical properties of samples collected in pasture, primary rainforest, and secondary rainforest in the State of Pará, Brazil. The mean values and results from an ANOVA with a post-hoc Tukey-Kramer test are tabulated. M.O. = organic matter. H+Al = total soil acidity, SB = sum of exchangeable bases (Ca + Mg + K), CTC = cation exchange capacity, m = aluminum saturation. mmolc = millimoles of charge,V = base saturation as a percentage of CTC, m = aluminum saturation as a percentage of CTC.

Suppl. Table 13. Soil physico-chemical properties of samples collected in pasture, primary rainforest, and secondary rainforest in the State of Rondônia, Brazil. The mean values and results from an ANOVA with a post-hoc Tukey-Kramer test are tabulated. M.O. = organic matter, H+Al = total soil acidity, SB = sum of exchangeable bases (Ca + Mg + K), CTC = cation exchange capacity. m = aluminum saturation, mmolc = millimoles of charge, V = base saturation as a percentage of CTC, m = aluminum saturation as a percentage of CTC.

