## Supplemental Methods for "Rainforest-to-pasture conversion stimulates soil methanogenesis across the Brazilian Amazon"

**Supplemental Materials and Methods for Kroeger et al. 2020**

**GPS Coordinates and Description**

In Pará, the sample site names are as follows: two primary rainforest sites are Pará-PF1 (S3° 17.736', W54° 57.776') and Pará-PF2 (S2° 51. 29', W54° 57.394'), pasture is Pará-P (S3° 17.614', W54° 51.046'), and secondary rainforest is Pará-SF (S3° 17.979', W54° 53.45'). The secondary rainforest is ~ 40 years old and the pasture (Pará-P) was established 23 years ago in 1996. In Rondônia, the sample site names are as follows: two primary rainforest sites are Rondônia-PF1 (S10º 8.435', W62º 54.000') and Rondônia-PF2 (S10º 8.54667', W62º 52.92000'), pasture is Rondônia-P (S10º 10.22167', W62º 49.95667'), and secondary rainforest is Rondônia-SF (S10º 9.63167', W62º 47.86500'). The secondary rainforest is 20 years old and the pasture is 47 years old.

**Quantitative PCR Reaction Mixtures and Thermocycler Protocols**

The particulate methane monooxygenase alpha subunit gene (*pmoA*) was amplified using the primer pair A189f/mb661r[34-35], and the gene for the methyl coenzyme M reductase alpha subunit (*mcrA*) was amplified using the primer pair mlas/mcra-rev[36]. The master mix to target *pmoA* consisted per reaction of 1.4 µl of PCR water, 1× KlenTaq Mutant Reaction Buffer, 0.2 mM each dNTP, 0.13 µM forward primer, 0.13 µM reverse primer, 1× EvaGreen Dye (Biotium, Fremont, CA), 200 ng/µL bovine serum albumin, 3 mM $MgCl_2$ , 0.2 µl of Omni KlenTaq polymerase (DNA Polymerase Technology, St. Louis, Missouri), and 10 µl of DNA template or PCR-grade water. The master mix to target *mcrA* was the same as above except for 10.4 µl of PCR-grade water and 1 µl of DNA template or PCR-grade water.

For the functional marker gene particulate methane monooxygenase alpha subunit (*pmoA*), we used the following thermocycler protocol: 10 min at 95°C, 40 cycles of 95°C for 15 s, 58°C for 30 s, 68°C for 45 s, and 82°C for 12 s followed by a plate read, incubation at 68°C for 5 min, melting curve from 65°C to 95°C with a read every 1°C and temperature holds of 1 s. For the functional gene marker methyl coenzyme M reductase alpha subunit (mcrA), we used the following thermocycler protocol: 20 min at 37°C, 5 min at 95°C, 40 cycles of 95°C for 30 s, 55°C for 45 s, 68°C for 30 s, and 83°C for 12 s followed by a plate read, incubation at 68°C for 7 min, melting curve from 65°C to 95°C with a read every 0.5°C and temperature holds for 1 s.

**Amplicon Sequencing**

The 3 genes of interest (16S v4 region, *pmoA*, and *mcrA*) were amplified using custom dual-indexed PCR primers designed by the University of Oregon Genomics & Cell Characterization Core Facility. These primers contain 4 major elements: the forward or reverse template binding sequence, an 8-nucleotide library barcode, either of the standard Illumina p5 or p7 adapter sequences, and a 3' phosphorothioate (PTA) modification to prevent 3'-5' exonuclease activity. For the 16S rRNA gene, primers 515F and 806R were used. For pmoA, the forward primer was AATGATACGGCGACCACCGAGATCTACACTATGGTAATTGTGGNGACTGGGACT*T*C*T*G*G and the reverse was CAAGCAGAAGACGGCATACGAGATAGTCAGTCAGCCCCGG MGCAACGTCYT*T*A*C*C. For mcrA, the forward primer was AATGATACGGCGACCACCGAGATC TACACTATGGTAATTGTGGTGGTGTMGGDTTCACMC*A* R*T*A and the reverse primer was

CAAGCAGAAGACGGCATACGAGATAGTCAGTCAGCCCGTTCATBGCGTAGTTVGGR T*A*G*T. A no-template control reaction was also included for each gene using Qiagen PCR-grade water (Qiagen, Hilden, Germany) in place of template gDNA.

The PCR reaction for all genes included the following for a total volume of 25 µL at 0.5 µM each primer: 12.5 µL of NEBNext® Q5® Hot Start HiFi PCR Master Mix (New England Biolabs, Ipswich, MA), 11.5 µL of combined forward and reverse PCR primer (each at 1.09 µM), and 1 µL of gDNA template. The PCR thermocycler protocol for 16S rRNA v4 region was 1 cycle at 98°C for 30 s, 25 cycles at 98°C for 10 s, 61°C for 20 s, 72°C for 20 s, 1 cycle at 72°C for 2 min. For the amplification of *pmoA* and *mcrA*, the PCR themocycler protocol was 1 cycle at 98°C for 30 s, 35 cycles at 98°C for 10 s, 65°C for 20 s, 72°C for 20 s, 1 cycle at 72°C for 2 min.

Following amplification, all libraries were purified with Omega MagBind TotalPure NGS (Omega Bio-tek Inc, Norcross, GA) via two subsequent 0.8 ratio bead cleanups to remove excess PCR primer. The purified libraries were quantified with Quant-iT™ high sensitivity dsDNA assay kit (Invitrogen, Carlsbad, CA) and characterized with the High Sensitivity NGS Fragment Kit on an Advanced Analytical Fragment Analyzer instrument (Agilent, Santa Clara, CA). Libraries were multiplexed to obtain equimolar representation (except for any libraries which quantified under 0.1 ng/uL and were therefore underrepresented in the pool).

The multiplexed pools for each of the 3 genes were combined in a final 1:1:1 ratio for sequencing on an Illumina MiSeq instrument using a v3 dual-indexed flow cell with PE 2x300 reads (Illumina, San Diego, CA). The sequencing library was spiked with 25% PhiX to increase nucleotide diversity. Custom sequencing primers for each of the 3 genes containing 3' PTA modifications were spiked into the reagent cartridge for Read 1, Read 2, and Index 1.

**Metagenome Library Prep and Sequencing**

Metagenome sequencing samples were prepared using the Nextera XT kit according to manufacturer's instructions (Illumina, San Diego, CA). Briefly, 1 ng of genomic DNA per sample was mixed with tagmentation buffer and enzyme and incubated for 5 minutes at 55°C. Samples were mixed with adapter sequences and amplified for 12 cycles with the following protocol: 1 cycle at 72°C for 3 min, 1 cycle at 95°C for 30 s, 12 cycles at 95°C for 10 s, 55°C for 30 s, 72°C for 30 s, 1 cycle at 72°C for 5 min. Samples were cleaned and size selected using Omega MagBind TotalPure NGS with a 0.9 bead ratio to remove excess PCR primer (Omega Bio-tek Inc, Norcross, GA). Libraries were characterized with the High Sensitivity NGS Fragment Kit on an Advanced Analytical Fragment Analyzer instrument (Agilent, Santa Clara, CA), and quantified using size corrected qPCR with KAPA library Quantification kit for NGS (Roche, Basel, Switzerland). Libraries were pooled at equimolar concentration into a single pool and run across two lanes of paired-end 150 sequencing on Illumina HiSeq 4000.

**Statistical Analysis of Sequences**

For Rondônia samples, Genbank annotations were rarefied to 4086593, 5673322, 4393131 for $CH_4$, $CO_2$, and NaAOc respectively. SEED Subsystem annotations were rarefied to 1718245, 2612838, 1942422 for $CH_4$, $CO_2$, and NaAOc respectively. For Pará samples, Genbank annotations were rarefied to 1280596, 1246399, 481062 for $CH_4$, $CO_2$, and NaAOc respectively. SEED Subsystems annotations were rarefied to 1503629, 1445579, 546595 for $CH_4$, $CO_2$, and NaAOc respectively.
