## Supplementary figures and images for "Rainforest-to-pasture conversion stimulates soil methanogenesis across the Brazilian Amazon"

### Supplemental Figures

SFigure 1

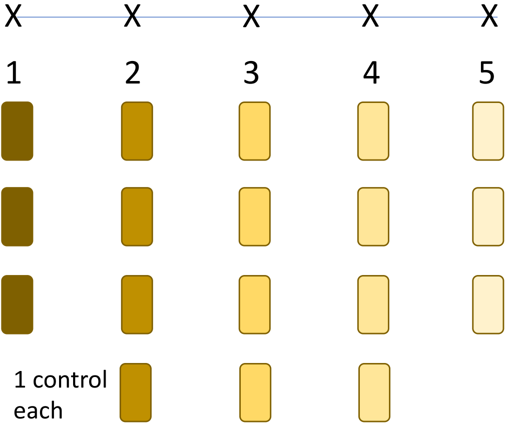


4 Land Uses for each Location:

- 2 Primary Rainforest
- 1 Pasture
- 1 Secondary Rainforest

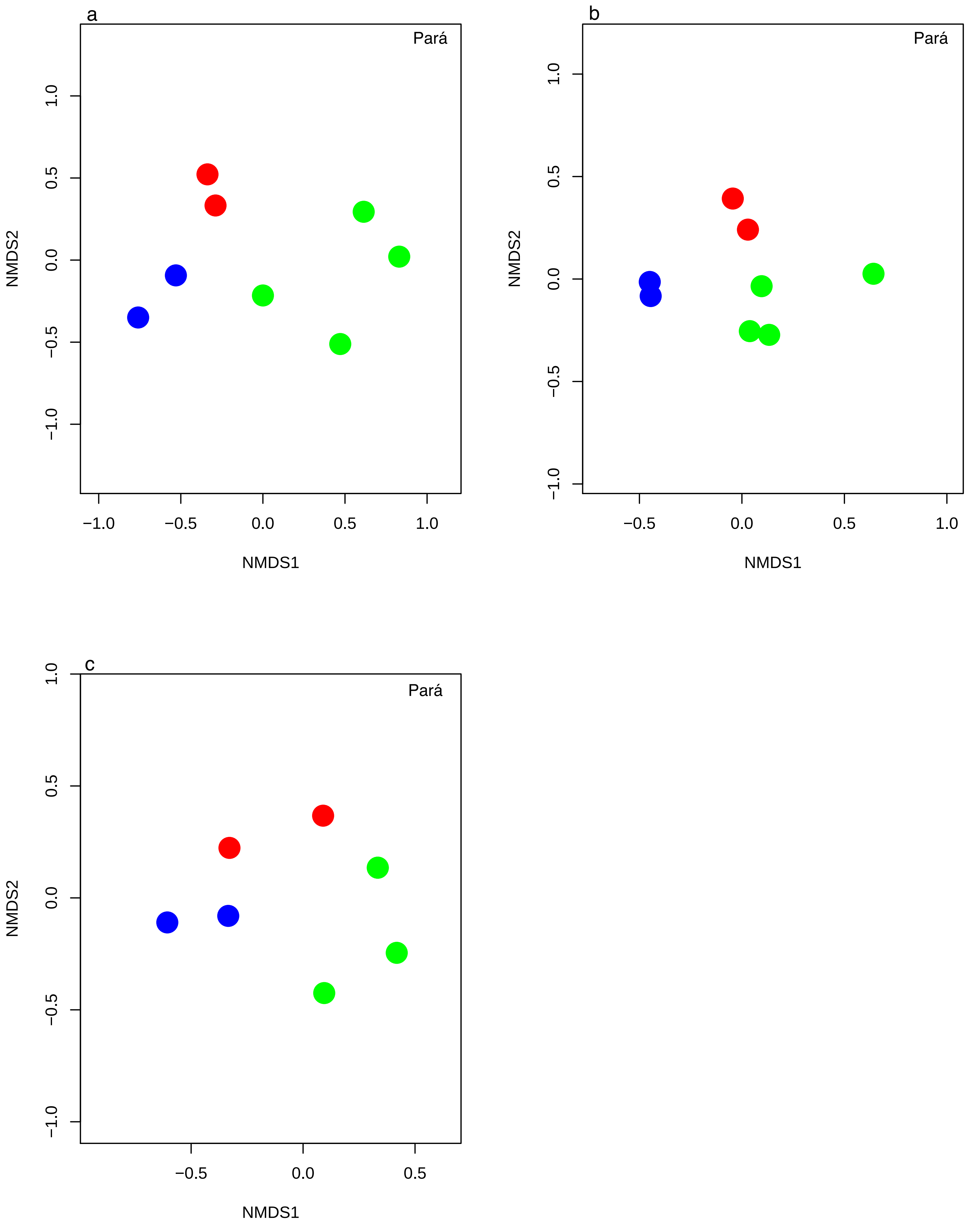

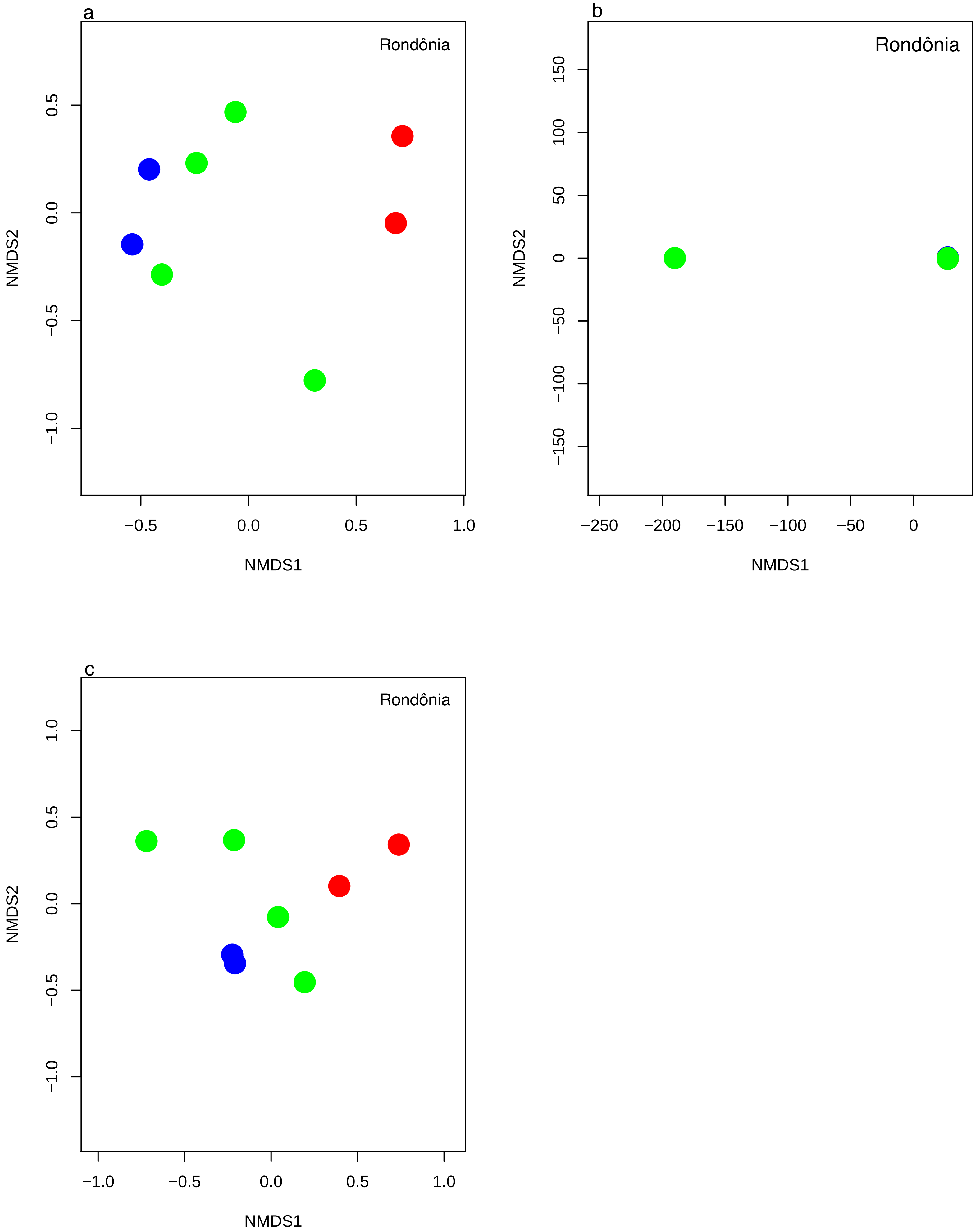

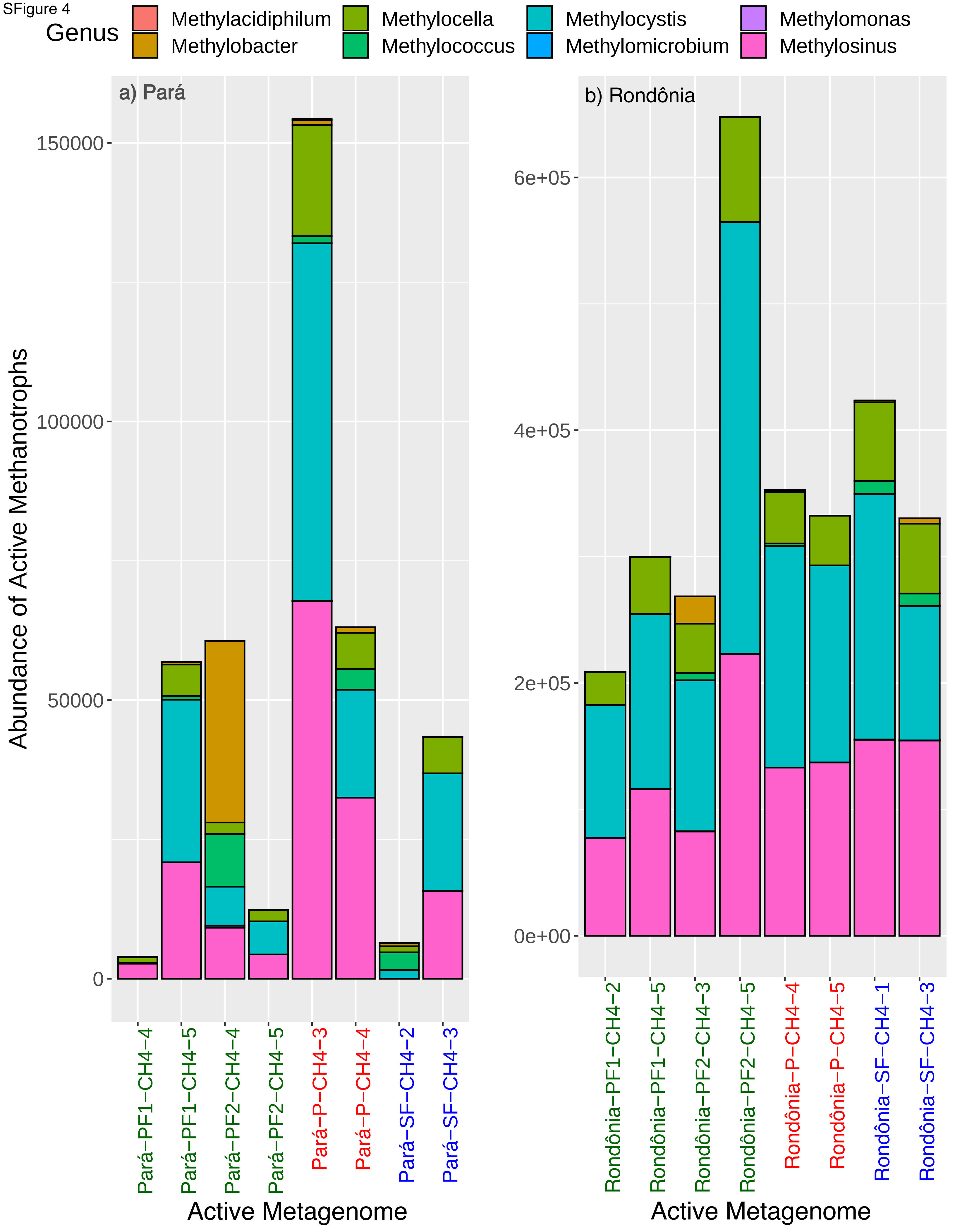

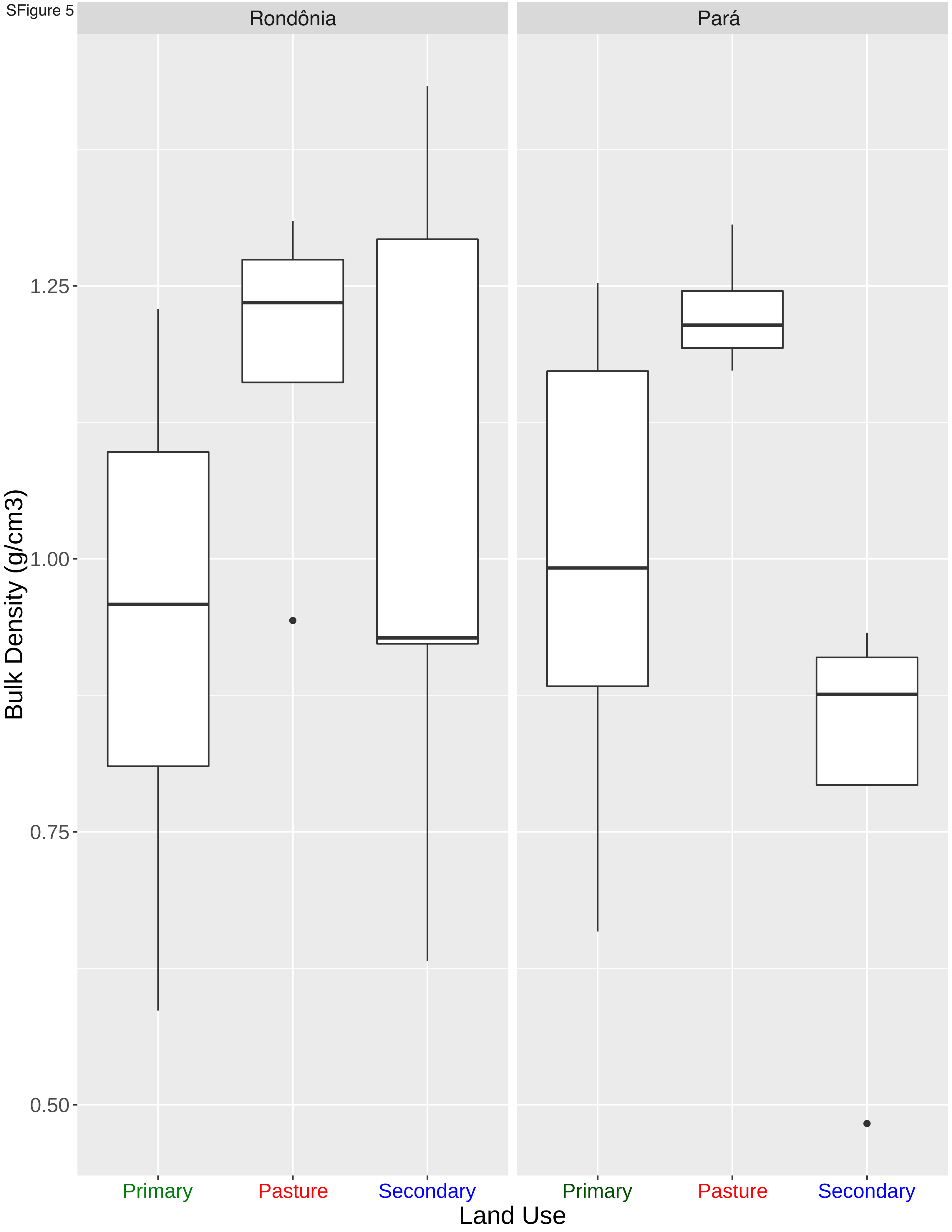

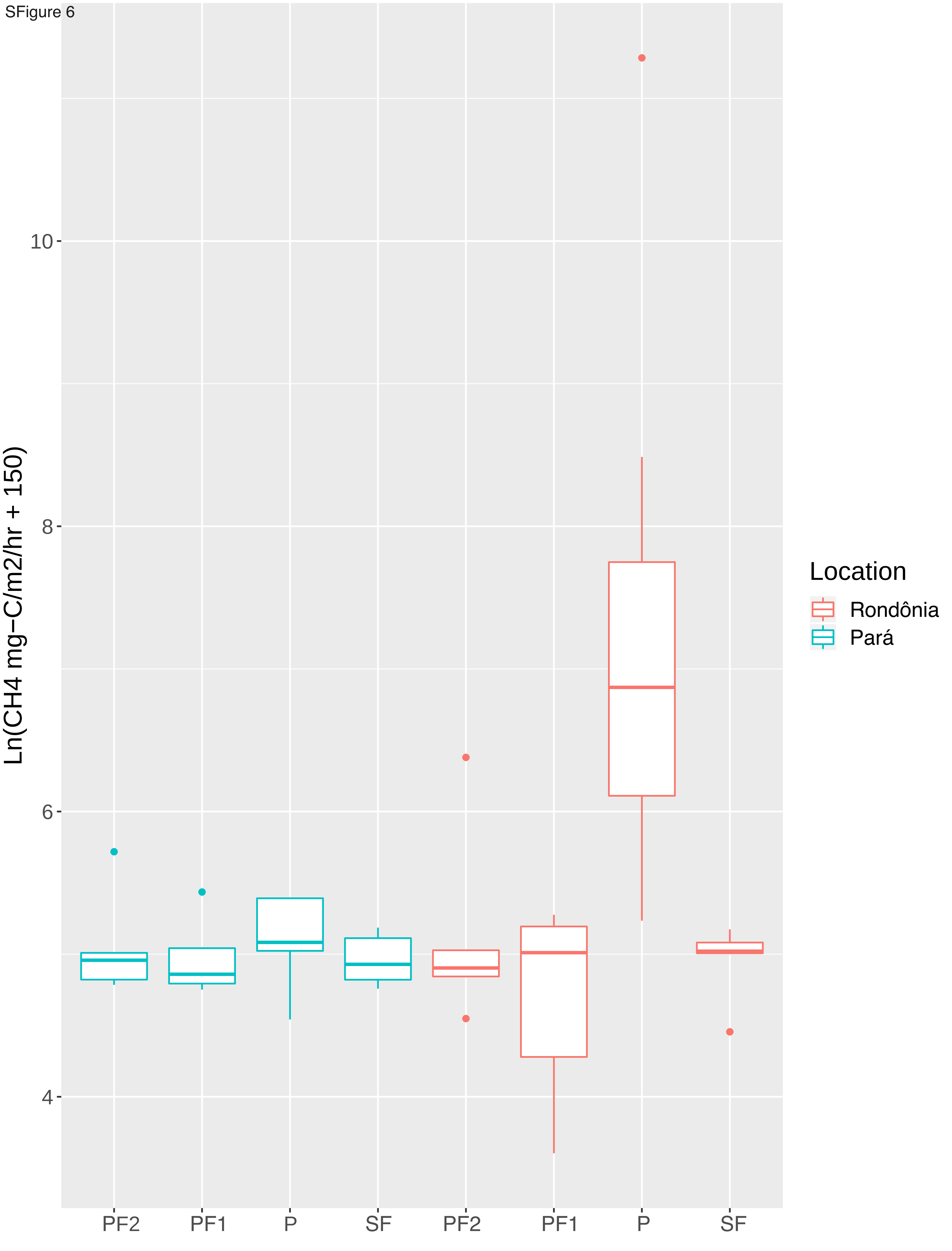

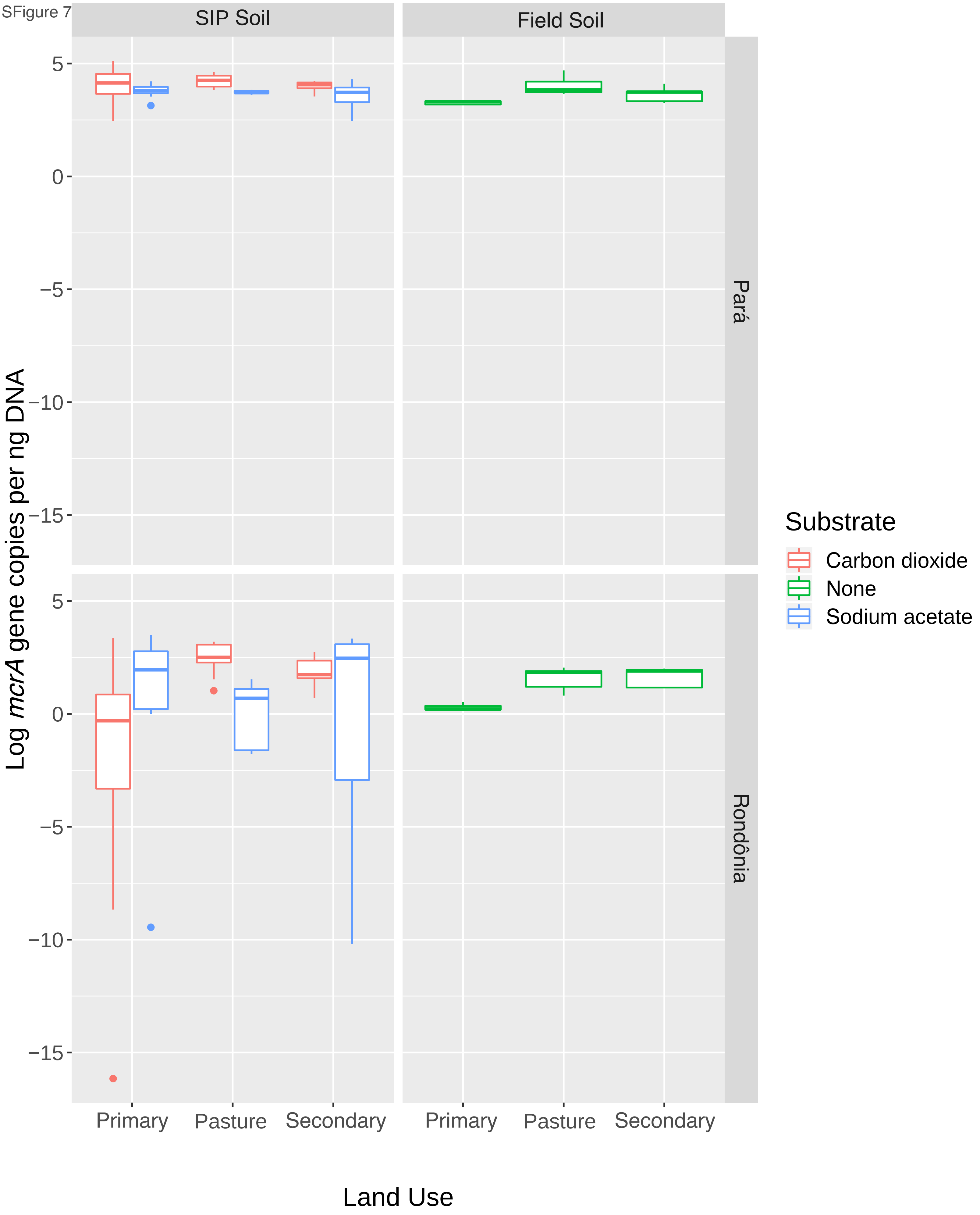

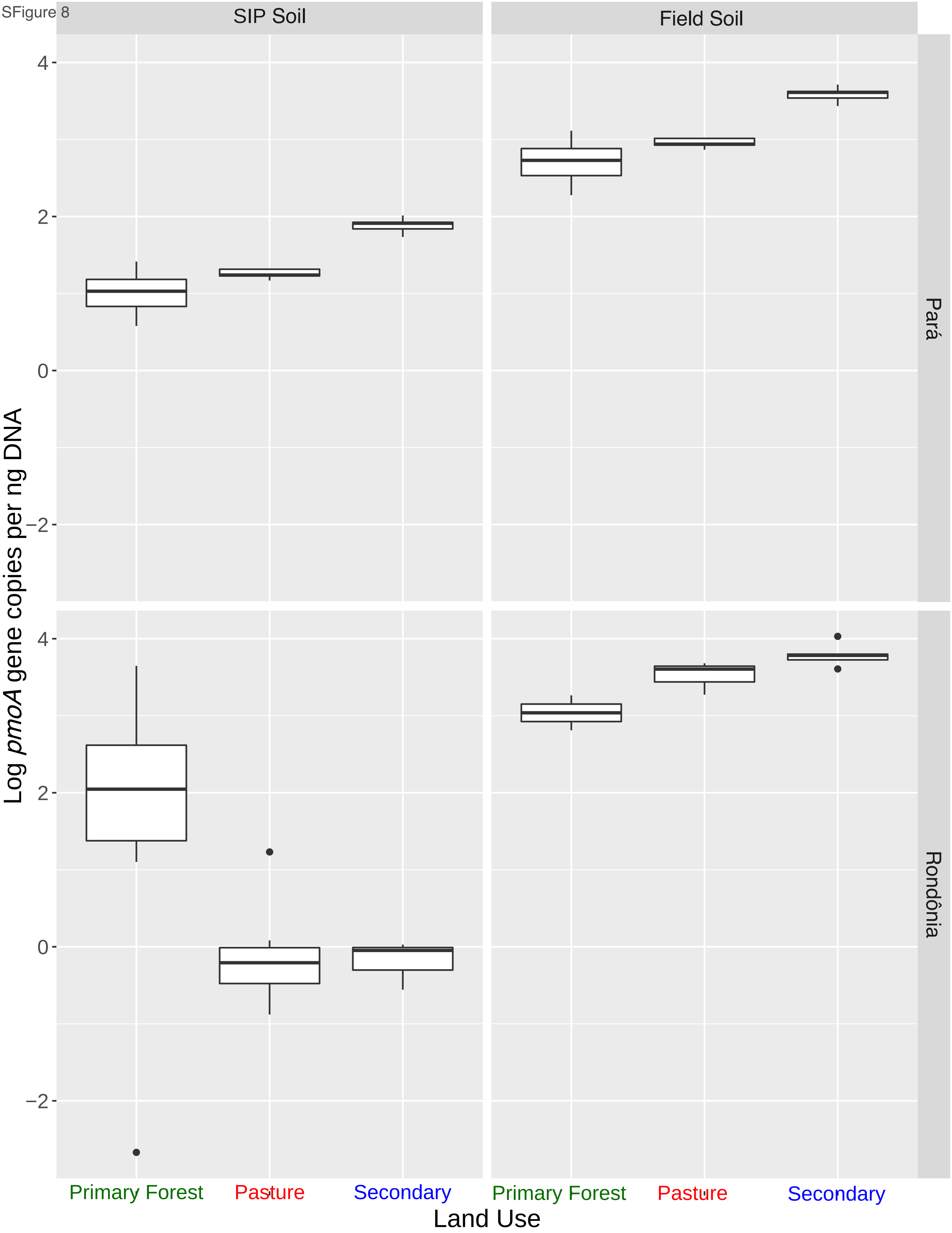
